# CRISPR/Cas9 ribonucleoprotein-mediated mutagenesis in *Sporisorium reilianum*

**DOI:** 10.1101/2023.10.17.562663

**Authors:** Janina Werner, Weiliang Zuo, Gunther Doehlemann

## Abstract

Clustered Regularly Interspaced Short Palindromic Repeats/CRISPR-associated protein 9 (CRISPR/Cas9) has become the state of art for mutagenesis in filamentous fungi. Here, we describe a RNP-mediated CRISPR/Cas9 for mutagenesis in *Sporisorium reilianum*. The efficiency of the method was tested *in vitro* with a cleavage assay as well as *in vivo* with a GFP-expressing *S. reilianum* strain. We applied this method to generate frameshift-, knock-in- and knock-out mutants in *S. reilianum* without a resistance marker by using an auto-replicating plasmid for selection. The RNP-mediated CRISPR/Cas9 increased the mutagenesis efficiency and firstly enables a marker-free genome editing in *S. reilianum*.

## 1. Introduction

The smut fungi consist of more than 1,500 species and are highly economically important due to their infection of relevant crops such as barley, sorghum, wheat and maize (Begerow *et al*., 2004). The majority of smut fungi infect their host systemically through the roots and replace the inflorescences by teliospores without causing symptoms during early infection stages (Martinez *et al*., 2002; Laurie *et al*., 2012). One example for this systemic infection is *Sporisorium reilianum f. sp. zeae*, which is the causal agent of maize head smut. *S. reilianum* is closely related to the intensively investigated model organism *U. maydis*. However, they differ in their mode of infection as well as in the site of the symptom development (Begerow *et al*., 2006; Steinberg *et al*., 2008). In 2010, a genome sequence of *S. reilianum f. sp. zeae* was published, which together with the *U. maydis* genome provided the foundation for systematic identification and genetic manipulation of effector genes contributing to virulence (Kämper *et al*., 2006; Schirawski *et al*., 2010). Genome comparison of *U. maydis* and *S. reilianum* revealed conserved effector genes even though they differ drastically in their pathogenesis on the same host, *Zea mays*. To characterize effector genes and their contribution to virulence, knock-out mutants are generated and compared to the wild type.

In the past, *U. maydis* knock-out mutants were generated using PCR-amplified donor templates with resistance markers for gene replacements (Brachmann *et al*., 2004). Importantly, it was shown that not only the genomic locus, but also the integration of resistance markers can negatively influence the expression of re-integrated genes (Schmitz *et al*., 2020). Recently, the mutagenesis of *U. maydis* was drastically improved with a marker-free approach using Clustered Regularly Interspaced Short Palindromic Repeats/CRISPR-associated protein 9 (CRISPR/Cas9) (Schuster *et al*., 2016; Zuo *et al*., 2020), and further developed for a seamless gene conversion approach (Zuo *et al*., 2021). In contrast to *U. maydis*, the generation of knock-out mutants in *S. reilianum* is still dependent on resistance markers and multiple gene knock-outs are hampered by the limited number of four available resistance markers (Brachmann *et al*., 2004). However, for *S. reilianum* the plasmid-based CRISPR/Cas9 transformation as used in *U. maydis* has not been successful.

CRISPR/Cas9 originating from the adaptive immune system of *Streptococcus pyogenes* has been broadly adapted to many eukaryotic systems. It’s a versatile tool for mutagenesis in various filamentous fungi (Schuster and Kahmann, 2019). The delivery strategies of CRISPR/Cas9 differ between fungal species: (i) stable genomic integration of *cas9*, (ii) transient delivery of Cas9 where the expression of Cas9 is dependent on selection pressure of a self-replicating plasmid or a telomere vector, respectively (Schuster *et al*., 2016; Leisen *et al*., 2020), or (iii) ribonucleoprotein complex (RNP)-mediated transformation (Foster *et al*., 2018; Leisen *et al*., 2020). Here, we describe CRISPR/Cas9 applications in *S. reilianum* using a RNP-mediated transformation approach. We demonstrate the generation of frameshifts as well as knock-out mutants mediated by RNPs and thereby generally improve the mutagenesis, and firstly enable a marker-free editing in *S. reilianum*.

## 2. Material

### 2.1 Equipment

a. PCR machine
b. Microfuge for PCR tubes
c. Tabletop centrifuge
d. 37°C incubator
e. 28°C incubator
f. Polyacrylamide gel electrophoresis (SDS-PAGE) equipment
g. Agarose gel electrophoresis equipment
h. Chemi-Doc MP System (or equivalent imaging system), with GFP filter
i. Geldoc: Visualization of DNA by UV radiation using a gel documentation unit (Peqlab/VWR, Radnor, USA)

### 2.2 Reagents and consumables

#### 2.2.1 sgRNA synthesis

a. T4 DNA Polymerase (NEB, Catalog No. M0203S)
b. NEB 2.1 buffer 1 (50 mM NaCl, 10 mM Tris-HCl, 10 mM MgCl2, 100 μg ml-1 BSA, pH 7.9)
c. dNTPs (DNA) (Roth, K039.1)
d. NucleoSpin® Gel and PCR clean-up (Machery and Nagel, Catalog No. 740.609.250)
e. HiScribe^®^ T7 High Yield RNA Synthesis Kit (NEB, Catalog No. E2040S)
f. DNase I (ThermoFisher, Catalog No. EN0521)
g. RNA Clean & Concentrator 25 kit (Zymo Research, Orange, CA, USA, Catalog No. R1017 & R1018)
h. Purple Loading Dye (NEB, Catalog No. B7024S; ingredients: 2.5% Ficoll®-400, 10 mM EDTA, 0.08% SDS, 0.02% Dye 1, 0.02% Dye 2, pH 8)
i. Nuclease-free water
j. PCR tubes and 1.5 ml Eppendorf tubes
k. 10x TBE buffer

#### 2.2.2 Formation of RNP and *in vitro* cleavage assay

a. EnGen^®^ Spy Cas9 NLS + NEBuffer r3.1 (<u>M0667</u>)
b. 500 mM Ethylenediaminetetraacetic acid (EDTA)
c. Proteinase K (ThermoFisher, Catalog No. EO0491)
d. 100 bp Ladder (NEB, Catalog No. N3231S)
e. 50x TAE buffer (2 M Tris base, 2 M acetic acid, 10% (v/v) 0.5 M EDTA (pH 8))
f. 1× TAE buffer (2%(v/v) 50x TAE buffer, 98% (v/v) VE water)
g. Agarose
h. 1% Ethidium bromide solution (Carl Roth)

#### 2.2.3 Protoplasting and transformation of *S. reilianum*

a. Novozym 234 (Novo Nordisc; Denmark)
b. SCS buffer (Solution 1: 0.6% (w/v) Natrium citrate, pH 5, 18.2% (w/v) Sorbitol, Solution 2: 0.4% (w/v) Citrate acid, 18.2% (w/v) Sorbitol; solution 1 and 2 are mixed until pH 5.8 is reached (approx.: 5:1 ratio from Solution 1 and 2; autoclaved)
c. STC buffer (50% (v/v) 2M Sorbitol, 1% (v/v) 1 M Tris-HCl, pH 7.5, 10% 1 M CaCl2, sterile-filtrated)
d. STC/40% PEG (60% (v/v) STC-buffer, 40% (w/v) PEG (MW 3350, Sigma P-3640); sterile-filtrated)
e. Regeneration agar light (1% (w/v) Yeast-Extract (Difco), 0.4% Bacto™ –Pepton (Difco), 0.4% (w/v) Sucrose (Roth), 18.22% Sorbitol (Sigma S-1876), 1.5% (w/v) Agar (Difco))
f. Potato dextrose agar (PDA)
g. Carboxin (5 mg/ml); (ThermoFisher, Catalog No 467620250)
h. Heparin (15 mg/ml)
i. Sterile cut tips (1000 μl and 20 μl)

### 2.3 Fungal strains

*S. reilianum* strains were stored at -80°C in 30% Glycerol. For transformation, *S. reilianum* wild type strains SRZ1 and SRZ2 (Schirawski *et al*., 2010) were used.

## 3. Method

### 3.1 *In vitro* transcription of sgRNA

a. Design protospacer in CHOPCHOP sgRNA designer (https://chopchop.cbu.uib.no/) using *Ustilago maydis* as target organism. Choose the protospacer sequence starting with a “G”, which is needed for initiating the transcription by T7 RNA Polymerase. If there is no desired protospacer starting with “G”, add an additional “G” upstream of the chosen protospacer sequence (21nt).
b. Add T7 RNA polymerase promoter sequence and <u>overlapping scaffold sequence</u> upstream and downstream of the chosen protospacer sequence, respectively, and order the gene-specific oligonucleotide (Table 1). In addition, a reverse complementary constant oligonucleotide is needed, which harbors the scaffold and terminator sequence and a 20 nt overlap to the scaffold sequence of the gene-specific oligonucleotide.
c. Mix both oligonucleotides in 1:1 ratio as following: 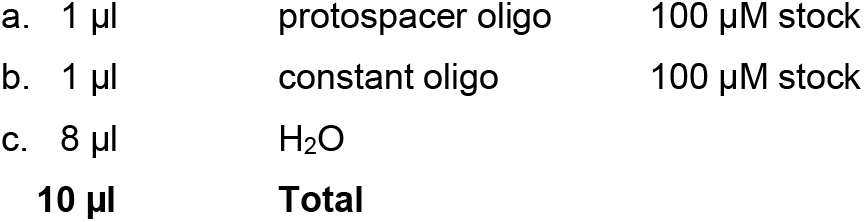 and anneal the oligos using the following program in PCR machine:

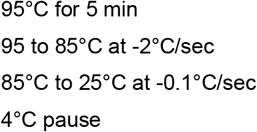
d. Add T4 DNA polymerase to fill in the overhangs:

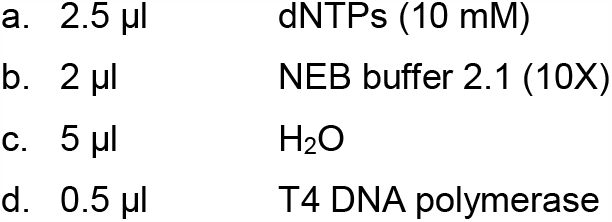
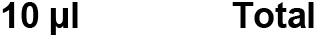 and incubate at 12°C for 20 min in a PCR machine.
e. Purify the product with PCR clean-up kit (Machery and Nagel), measure the concentration by Nanodrop and verify the PCR product on a 2-3% TAE agarose gel.
f. Use 2 μg of the resulted DNA from above as template and the HiScribe T7 High Yield RNA Synthesis Kit for the following reaction (NEB, protocol for small RNAs):

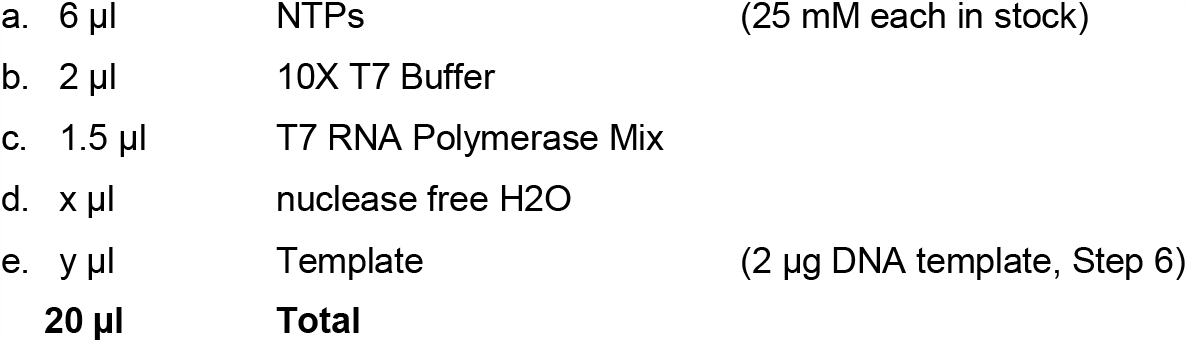 Flip the tube, vortex shortly and incubate at 37 °C overnight.
g. Next day, add 14 μl Nuclease-free H_2_0, 4 μl DNase I buffer (10 X), 2 μl DNase and incubate at 37°C for 15 min.
h. Purify the resulted sgRNA with the RNA Clean & Concentrator 25 kit (Zymo, Catalog No R1017 & R1018) and use the manufacturer’s protocol (manual, page 5). (Optional: Check quality of the RNA on 10% denaturing PAA gel using TBE buffer (89 mM Tris base, 89 mM boric acid, 2 mM sodium EDTA) and 8 M urea, and TBE as running buffer (Leisen *et al*., 2020)).

**Table 1:**
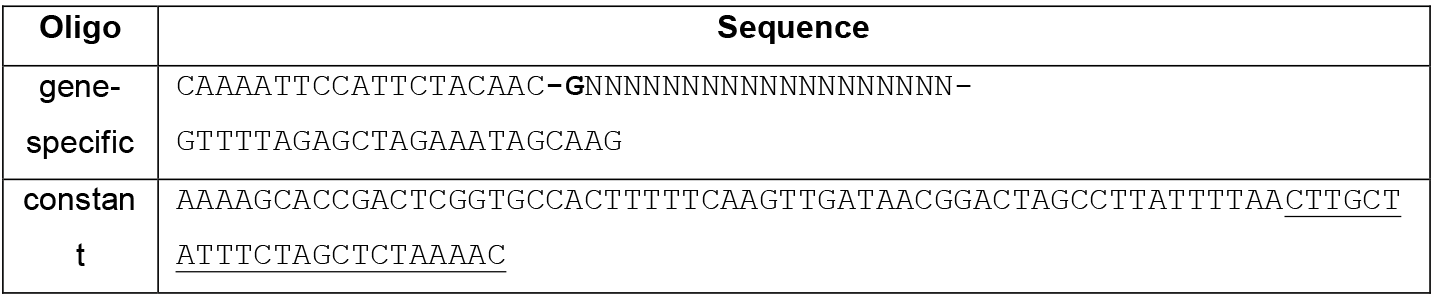
Sequences of Oligonucleotides for sgRNA synthesis.

### 3.2 *In vitro* cleavage assay

a. To test the *in vitro* efficiency of the designed sgRNA, Cas9 is mixed with the sgRNA in 1:1 molar ratio (15 μM) and incubated for 10 min at RT (Figure 1B).
b. Afterwards, 333 ng of a DNA cleavage template (PCR product of the region of interest) is added (Figure 1C).
c. After 1h, 2h and 3h 10 μl samples are taken (Figure 1D) and the reaction is stopped by the addition of 1 μl 500 Mm EDTA and 1 μl Proteinase K. Subsequently, the reaction is incubated for 30 min at 50°C for degradation of Cas9.
d. After the collection of all samples, cleavage is checked on an 1.5% agarose gel (Figure 1E).

**Figure 1:**
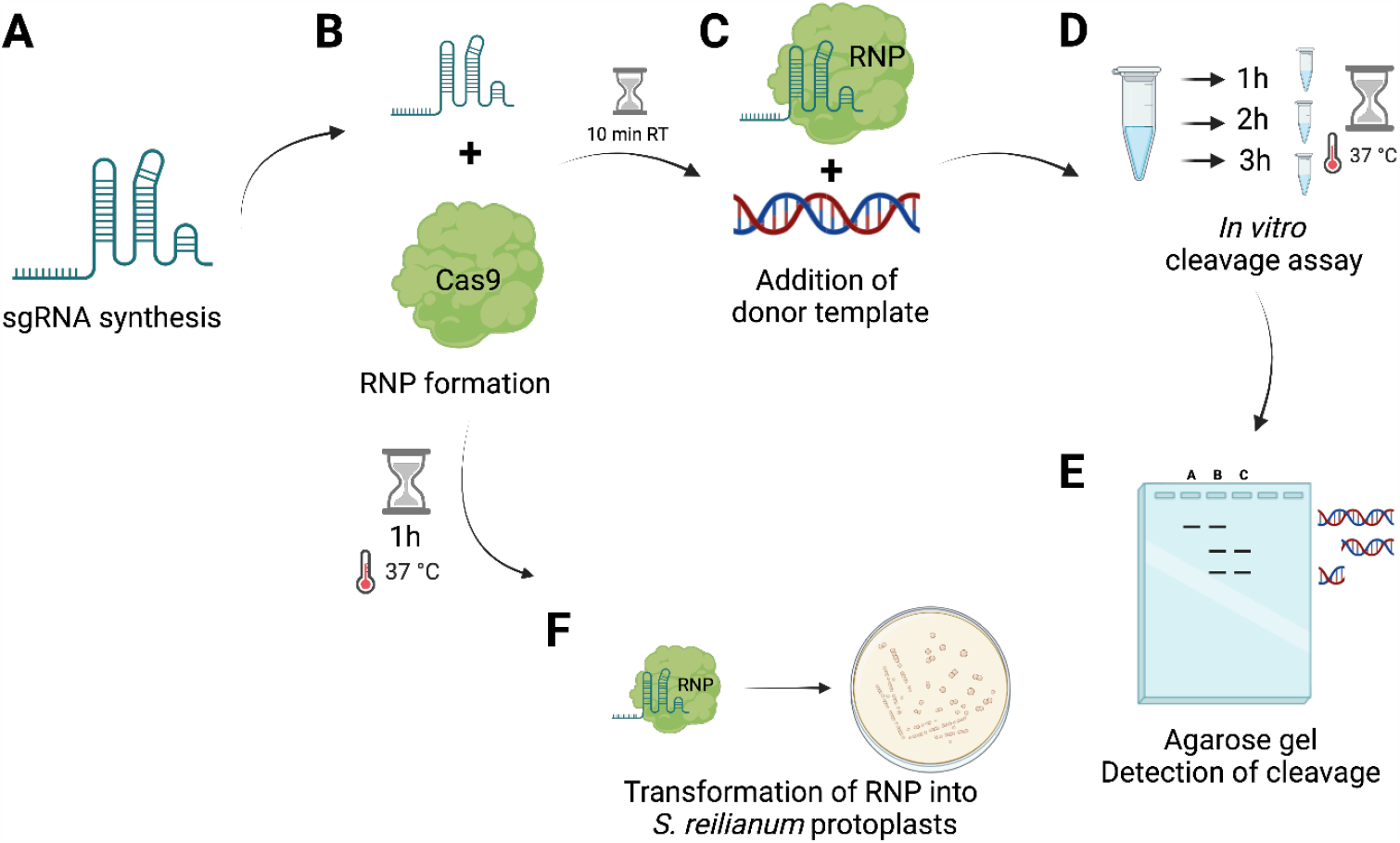
Workflow of RNP-mediated transformation in *S. reilianum*. **(A)** *In vitro* synthesis of sgRNA using T7 HiScribe kit. **(B)** RNP formation of *in vitro* transcribed sgRNA with *Sp*Cas9. **(C)** Alternatively: To perform an *in vitro* cleavage assay, incubation at RT for 10 min and subsequently the addition of a donor template (Amplification of the gene of interest region) and incubation at 37°C for 3h is conducted. **(D)** Sampling of 10 μl reaction mix after 1h, 2h and 3h (or alternatively overnight). **(E)** Visualization of *in vitro* cleavage on a 1.5% agarose gel. **(F)** RNP incubation for 1 h at 37°C prior transformation into *S. reilianum* protoplasts. Figure: biorender.com.

### 3.3 Assembly of RNP for transformation into *S. reilianum*

2 μg of the *in vitro* transcribed sgRNA targeting the gene of interest is used and mixed with 6 μg of *Sp*Cas9. Subsequently, r3.1 buffer of NEB and water are added in a minimum volume (Figure 1B). After mixing and centrifugation, the reaction is incubated for 1h at 37 °C prior transformation (Figure 1B).

### 3.4 Transformation of *S. reilianum*

*S. reilianum* protoplasts are prepared as described previously (Brachmann *et al*., 2004). For RNP transformation, protoplasts are thawn on ice for 5 min, and then a self-replicating plasmid with antibiotic resistance cassette (pNEBUC - Carboxin; Brachmann *et al*., 2004), the RNP, 1 μl of 15 mg/ml Heparin and optionally 1.5 μg of a donor template is added (Figure 2). After a 10 min incubation on ice, 500 μl of STC/PEG is added and re-suspended carefully with a tip-cut blue tip until it looks homogenous (5-8 times pipetting up and down). Followed by an incubation for 15 min on ice, the protoplasts are spread on a regeneration agar light plate with 2 layers (bottom layer: corresponding selective antibiotic (for pNEBUC – carboxin), top layer: without antibiotic resistance). After 4 days, transformants are singled out with a blue tip on PDA+Carboxin (2.5 μg/ml) for 2-3 days. Afterwards, a single colony is transferred for 2 days to PDA plates to lose the resistance. Subsequently, DNA is isolated (Hoffman and Winston *et al*., 1987) and used for further confirmation (see 3.6). The efficiency of the RNP CRISPR/Cas9 can for instance be tested with GFP fluorescence as a readout (Figure 3A). To test the efficiency in *S. reilianum*, a strain harbouring a single integration of GFP controlled by pOTEF (constitutive promoter) was generated in the *ip* locus of SRZ2 strain and confirmed via Southern blot. For the transformation of *S. reilianum* protoplasts a sgRNA against GFP together with the Cas9 in a RNP (see 3.3) and an auto-replicating plasmid (pNEBUC) for selection on regeneration agar was used. Transformants were singled out after 4 days of incubation at 28°C on PDA+Cbx (2.5 mg/ml) and after 2 days transferred to PDA plates and checked for their fluorescence using a Chemidoc.

**Figure 2:**
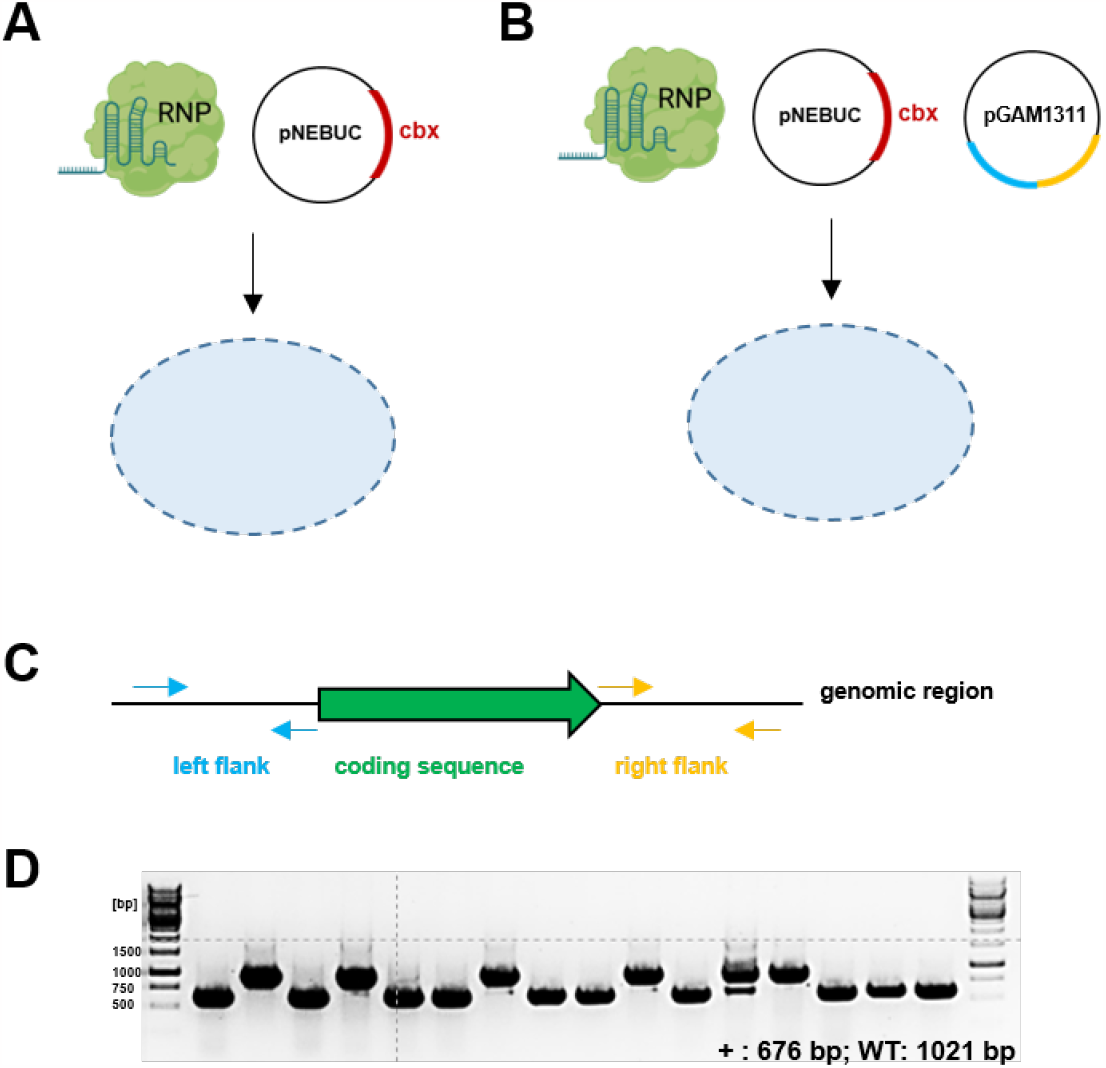
Transformation of *S. reilianum* using RNP with and without a donor template. **(A)** Generation of frameshift mutants using RNP-mediated transformation. **(B)** Generation of knock-out mutants using RNP-assisted homologous recombination with a donor template. **(C)** Donor template design for the generation of a knockout. 1kb flanking regions of the coding sequence are amplified by PCR with overhangs for Gibson assembly into backbone TK#1_pAGM1311 (MOCLO backbone, Level -1). **(D)** Example of deletion mutant verification PCR using LF forward primer and RF reverse primer (expected sizes: WT: 1021 bp, mutant (+): 676 bp; efficiency: 62.5% (10/16)). The efficiency can vary between different genomic loci. Figure 2A+B: biorender.com.

**Figure 3:**
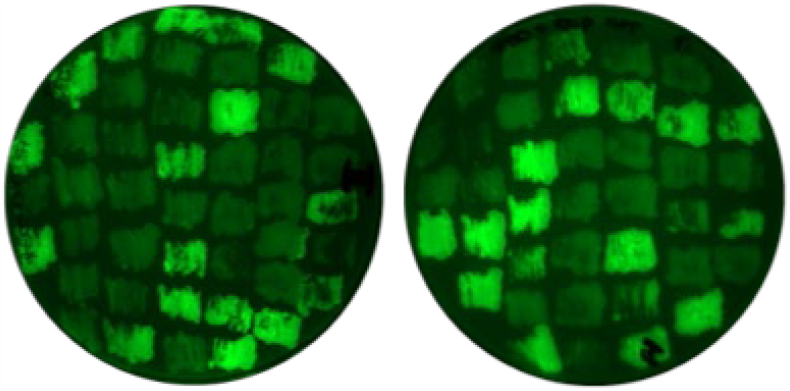
GFP as target for RNP-mediated transformation in *S. reilianum*. An example shows the efficiency of RNP-mediated CRISPR/Cas9 transformation in *S. reilianum*. A *S. reilianum* SRZ2 strain expressing GFP under pOTEF promoter was generated. sgRNA+Cas9 targeting GFP coding sequence was transformed, and mutants with frameshift lose the GFP signal. Efficiency for GFP sgRNA: ∼ 41 % (34/83).

### 3.5 RNP-assisted homologous recombination to generate a knock-out in *S. reilianum*

For the generation of an antibiotic resistance free knock-out mutant in *S. reilianum*, a CRISPR/Cas9 mediated homology-directed repair is exploit. To do this, a donor template is generated by cloning the 1 kb homology flanking regions of the target gene into a MOCLO vector TK#1_pAGM1311 by Gibson assembly (Figure 2C). For the transformation of *S. reilianum* protoplasts (see 3.4), the donor template together with an self-replicating plasmid (pNEBUC) and the RNP with a sgRNA against the target region was used. The transformants were transferred as described above (see 3.4). A PCR was conducted with the forward primer of the left flank and the reverse primer of the right flank (Figure 2C) and compared to the wild type (Figure 2D). Putative positive mutants from PCR are selected for further verification via Southern blot (Southern, 1975; Sambrock *et al*., 1989) using the deletion construct (left flank + right flank), previously used as a donor template, as probe for hybridization.

## Acknowledgements

This project has received funding from the European Research Council (ERC) under the European Union’s Horizon 2020 research and innovation programme (grant agreement No 771035), as well as funding by the Deutsche Forschungsgemeinschaft (DFG, German Research Foundation) under Germany’
ss Excellence Strategy-EXC-2048/1-Project ID: 390686111

